# Therapeutic targeting of MYBPC3 mutation-specific hypertrophic cardiomyopathy guided by network modeling

**DOI:** 10.64898/2026.09.03.749173

**Authors:** Pichayathida Luanpaisanon, Caitlin M. Pavelec, Leigh A. Bradley, Jeffrey J. Saucerman, Matthew J. Wolf

**Affiliations:** Department of Biomedical Engineering University of Virginia, Charlottesville VA, USA; Division of Cardiovascular Medicine, University of Virginia, Charlottesville VA, USA; Robert M. Berne Cardiovascular Research Center, University of Virginia, Charlottesville VA, USA

## Abstract

Hypertrophic cardiomyopathy (HCM) is a leading cause of sudden cardiac death with genotype positive cases most commonly associated with mutations of cardiac myosin binding protein-C (MYBPC3). Recently approved drugs for HCM target the myofilaments rather than the aberrant molecular signaling pathways that drive long-term remodeling. Here, we identified a patient with familial HCM that was associated with a MYBPC3 W1078 truncation mutation. A CRISPR knock-in mouse model of the orthologous mutation MYBPC3 W1082 exhibited marked cardiac hypertrophy including wall thickening, reduced ejection fraction, and decreased survival. To identify pathways, mechanisms, and potential candidate therapeutics, we integrated the MYBPC3 mutation into a computational network model of the signaling underlying familial cardiomyopathy. The network model predicted that the MYBPC3 mutation drove hypertrophy through mTOR/PI3K pathways, consistent with the results of RNA sequencing of cardiomyocytes of MYBPC3^W1082*/W1082*^ mice. A virtual drug screen using FDA-approved drugs predicted that the mTOR inhibitor, Rapamycin, could mitigate mutation-induced hypertrophy. We then experimentally validated the effects of Rapamycin on hypertrophic responses using cultured cardiomyocytes. Further, Rapamycin attenuated cardiac hypertrophy and fibrosis of MYBPC3^W1082*/W1082*^ mice *in vivo*. mTOR inhibitors (rapamycin and everolimus) were associated with a decreased incidence of cardiac hypertrophy associated diagnostic codes in patients in the FDA Adverse Events Reporting System. Query of electronic health records and echocardiograms from a University of Virginia cohort of patients treated with mTOR or calcineurin inhibitors showed that patients prescribed everolimus or tacrolimus had reduced LV wall thicknesses. Together, these studies suggest that targeting mTOR as a translationally relevant target for a mutation-induced hypertrophic cardiomyopathy, as well as demonstrating the utility of guiding precision therapies by iterating between network models and experimental validation.

## Key findings

- MYBPC3 W1082 mice recapitulate aspects of human HCM
- MYBPC3-specific HCM mutation network model predicts PI3K/AKT and mTOR pathways as key regulator of cardiac mass
- Myosin activator induces cardiomyocyte hypertrophy, mitigated by rapamycin
- Rapamycin reduce cardiac mass in MYBPC3 mutant mice

## Introduction

Hypertrophic cardiomyopathy (HCM) is the most prevalent inherited cardiac disease, affecting approximately 1 in 200–500 individuals worldwide and representing a major cause of sudden cardiac death in young people[1], [2]. HCM is defined by the presence of left ventricular hypertrophy (LVH) not secondary to causes such as hypertension, valvular disease, or infiltrative cardiomyopathy [3]. Despite a preserved ejection fraction (EF), patients frequently suffer debilitating symptoms including exertional dyspnea, chest pain, and syncope, driven by dynamic LV outflow tract obstruction, diastolic dysfunction, and microvascular ischemia [4], [5]. The disease course is highly variable: many individuals remain asymptomatic for decades, while others progress to heart failure, develop atrial fibrillation, or suffer life-threatening ventricular arrhythmias [6]. HCM is often caused by autosomal dominant mutations in genes encoding sarcomeric proteins, with over 1,500 distinct variants across at least 11 causal genes [7], [8]. MYBPC3 mutations are the single most frequent genetic cause of HCM, comprising roughly 40– 50% of pathogenic variants and predominantly consisting of mutations that result in premature truncation of the cMyBP-C protein and ultimately haploinsufficiency at the protein level [9], [10].

At the molecular level, loss of functional cMyBP-C disrupts its normally inhibitory influence on actomyosin cross-bridge cycling, leading to hypercontractility and increased ATP consumption [11], [12]. This elevated energetic demand triggers a cascade of downstream signaling events, including activation of calcineurin-NFAT, PI3K/AKT/mTOR, and MAPK/ERK pathways, that collectively drive pathological cardiomyocyte hypertrophy, sarcomeric disarray, and interstitial fibrosis resulting in the clinical manifestations of HCM [13], [14], [15], [16], [17]. Current management of HCM remains largely symptom-directed rather than disease-modifying. Beta-blockers, calcium channel blockers, and disopyramide are used to reduce LV outflow obstruction and improve diastolic filling, while implantable cardioverter-defibrillators are deployed for primary prevention of sudden death in high-risk patients [18]. In 2022, mavacamten, a selective allosteric inhibitor of β-cardiac myosin ATPase, became the first FDA-approved disease-specific pharmacological therapy for obstructive HCM, demonstrating significant reductions in LV outflow tract gradient and regression of LVH in clinical trials [19], [20]. However, mavacamten has important limitations including a major risk of excessive reduction in LVEF, requiring close echocardiographic monitoring and dose titration [21]. More fundamentally, mavacamten targets contractile mechanics rather than the upstream intracellular signaling networks that drive hypertrophic remodeling. Therefore, there remains a critical and largely unmet need for therapeutic strategies that directly target the signaling pathways responsible for pathological hypertrophy.

Mouse models have been indispensable tools for investigating HCM pathogenesis, however translation has been limited. Early transgenic models overexpressing truncated MYBPC3 constructs rely on non-physiological gene dosage, promoter-driven overexpression, and multi-copy integrations that confound interpretation of mutation-specific effects [22], [23]. Complete knockout models, including the widely used Mybpc3-null mouse, develop an aggressive dilated cardiomyopathy phenotype that poorly mirrors the clinical spectrum MYBPC3-mutation induced HCM in humans [24], [25]. Knock-in models carrying human HCM-associated variants offer greater fidelity, but most published knock-in lines have introduced variants distinct from those observed in human patients [26]. Furthermore, many preclinical studies have been conducted in isolation, focusing on a single phenotypic endpoint without integrating the multiscale signaling context necessary to identify actionable therapeutic targets. Systems biology approaches offer a powerful complement to experimental strategies by enabling comprehensive, network-level interrogation of disease mechanisms across multiple scales [27]. Logic-based ordinary differential equation models of cardiac signaling networks have previously been shown to accurately predict experimental outcomes across diverse contexts, including HCM and dilated cardiomyopathy (DCM), with the ability to simulate the effects of combinatorial pathway perturbations that would be impractical to test experimentally [28], [29].

Here, we report an integrated systems-level investigation of MYBPC3-associated HCM combining a clinically anchored mouse model, computational network modeling, quantitative transcriptomics, and pharmacological intervention. We generated a CRISPR-engineered mouse carrying the patient-specific MYBPC3^W1082*/W1082*^ variant, the murine ortholog of a pathogenic MYBPC3^W1078*^ truncation identified in an HCM patient. We then expanded a validated cardiac signaling network model to incorporate MYBPC3 loss-of-function, performed a virtual knockdown screen to identify the signaling pathways with greatest influence on cardiomyocyte hypertrophy, and integrated predictions with the DrugBank, FDA-approved drug-target database to prioritize therapeutic candidates. Transcriptomic profiling of MYBPC3^W1082*/W1082*^ mouse hearts provided orthogonal validation of computational predictions, and targeted in vivo and in vitro pharmacological experiments confirmed the efficacy of rapamycin, an FDA-approved mTOR inhibitor, in attenuating MYBPC3-driven hypertrophy. Finally, retrospective analysis of FDA adverse event reporting data and electronic health records at the University of Virginia provided population-level clinical corroboration. Together, these findings establish mTOR signaling as a mechanistically and therapeutically relevant in MYBPC3-associated HCM and demonstrate the power of a tightly integrated computational-experimental strategy for mutation-specific drug discovery.

## Materials and Methods

### Mice

All animal procedures were conducted in accordance with the University of Virginia Animal Care and Use Committee (UVA ACUC) Policy on Rodent Surgery and Perioperative Care under an approved protocol (Wolf Laboratory protocol no. 4080). The MYBPC3^W1082*^ mouse model was generated based on a hypertrophic cardiomyopathy pedigree identified at the University of Virginia Medical Center harboring a pathogenic MYBPC3^W1078*^ variant. The murine tryptophan residue at position 1082 is orthologous to human MYBPC3 tryptophan 1078.

CRISPR/Cas9-mediated genome editing was performed in the B6SJLF1 background by the University of Virginia Genome Editing and Mouse Modeling (GEMM) Core. A single guide RNA (sgRNA) targeting the Mybpc3 locus (5′-TGAGACTTGGGGTTTCAATGTGG-3′) was used in conjunction with a single-stranded DNA repair template (sequence provided below) designed to introduce a premature stop codon (TGA) in place of the native tryptophan codon (TGG). This substitution results in a truncating MYBPC3^W1082*^ allele, modeling the corresponding human disease-associated variant. The repair template sequence used is: *GCTTCTGCTCGGTCACTTCAGAACGCTCAGTGTGCCAATCAGATCTGTCTCCTGCAGACAAGCCAAGTCCTCCCCAGGATATCCGGATCGTTGAGACTTGAGGTTTCAATGTaaCTCTGGAGTGGAAGCCACCCCAAGATGATGGCAATACAGAGATCTGGGGTTATACTGTACAGAAAGCTGACAAGAAGACCATGGTG*

### Echocardiography

Six-to forty-week-old mice were anesthetized using isoflurane. The chest was exposed after hair removal, and a four-lead electrocardiogram (ECG) was collected during the procedure. Images were collected using a VevoView 1100 system. Images were collected in the long-axis (B-mode) and short axis (M-mode) on mice serially throughout the experimental period. Serial collection began prior to drug treatment or at 8 weeks of age whichever occurred first and was collected every two weeks during drug treatment progression or monthly for serial survival studies. Images were analyzed using Vevo Lab. Left-ventricle (LV) end-diastolic, end-systolic volume, and wall thickness were assessed, and ejection fraction calculated.

### Rapamycin Administration

Mice were given baseline (pre-treatment) echocardiography. Rapamycin (Selleck Chem-S1039) or vehicle control (10% Ethanol in PBS) was administered at 2mg/kg/day by intraperitoneal (i.p.) injection daily for 28 days. Echocardiography was collected at 2 and 4 weeks after drug administration had begun. At the end of the 28 days, mice were euthanized, and organs were harvested and weighed.

### Histological analysis

Tissues were fixed in 10% formalin for four hours and transferred to 70% ethanol prior to paraffin embedding. Paraffin-embedded tissues were cut into 10-micron sections, deparaffinized, and stained either with Masson’s Trichrome according to standard protocols or stained with a wheat germ agglutinin (WGA) antibody after antigen retrieval as previously described [30]. Masson’s trichrome imaging was performed was taken on a Leica LAS X Multi Channel Acquisition, while WGA imaging was performed on an Olympus Fluoview 1000 and are representative images of composite z-stacks. Images were taken with a 20X objective. Masson’s Trichrome staining was analyzed using FIJI with the color deconvolution macro (*340*), set to Trichrome [31]. The green channel was used for quantification of fibrotic area. A minimum of five Masson’s Trichrome images were taken per mouse, and fibrotic area was averaged across all images.

### Quantification of Cardiomyocyte cross-sectional area

Hearts were prepared as indicated in histological analysis including Sudan Black staining to quench autofluroesence. Sections were stained with DAPI (1:500 - Thermo Fisher D3571) and Wheat-Germ-Agglutinin-Alexa Fluor 647 (5 μg/m – Thermo Fisher W32466) during incubation. Sections were imaged on an Olympus Fluoview 1000 confocal microscope with five images per mouse taken in the left ventricle across three sections. 60 round nucleated cardiomyocytes per mouse were outlined across the images and were analyzed in ImageJ.

### Expanding Signaling Network Model with MYBPC3 HCM Mutation

Initially, we used a previously published logic-based differential equation model of the familial cardiomyopathy signaling network, composed of 39 nodes (proteins or mRNAs) and 82 reactions (edges), validated against 160 qualitative experimental data points curated from the literature [29]. This model correctly predicted 90% accuracy in HCM context versus 75% accuracy in DCM context, with an overall prediction accuracy of 83.8% [29]. We then expanded this network model to incorporate MYBPC3, its regulation, and the consequences of MYBPC3 gene mutation on the network [11], [12]. Specifically, we modeled the impact of MYBPC3 phosphorylation by PKA, PKC, PKD, and CaMKII to decrease myofilament Ca2+ sensitivity [32], as well as the impact of MYBPC3 gene mutation to de-repress myofilament Ca^2+^ sensitivity [33], [34]. The computer code for this expanded model is available under an MIT open-source license at: https://github.com/saucermanlab/MYBPC3network.

### Virtual Knockdown Screens to Identify Drug Candidates

As with past logic-based network models, we identified the most influential pathways by simulating individual knockdowns for each node in the network model [29], [35]. The steady-state activity of the model without simulation was obtained as basal conditions. We then knocked down the activity of each node one at a time (setting Ymax = 0) and subtracted the basal activity values from the values in the knockdown state to calculate changes in activity. Influence was measured and ranked by the highest change in activity following the knockout of the perturbed node. After the influential nodes (pathways) were identified, we matched protein targets of those nodes using the drug-target database at DrugBank [36] and obtained drug candidates predicted to have the greatest effects on cardiac Mass and Eccentricity.

### Experimental Validation in Cardiomyocytes

We isolated neonatal cardiomyocytes using the NeoMyts kit from Cellutron as described previously [37]. Cardiomyocytes were cultured with serum for 24 h in 96-well microplates, followed by a 16 h serum starve. Cardiomyocytes were then treated with one of two hypertrophic stimuli (10 μM phenylephrine, 10μM Omecamtive Mercarbil), 10% FBS (positive control), or serum-free media alone (negative control). At the same time, cells were treated with Rapamycin (Cat#J62473.MF) for 48 h.

To prepare for immunofluorescent imaging, cardiomyocytes were first fixed with 4% paraformaldehyde for 20 min and then permeabilized with 0.1% Triton-X for 15 min. Cardiomyocytes were blocked with 1% bovine serum albumin in PBS for 1 h, then treated with mouse anti-α-actinin primary antibody (Sigma-Aldrich Cat#A7811, RRID:AB_476766) at a concentration of 1:200 overnight. Cardiomyocytes were blocked with 5% goat serum in PBS for 1 h, then Alexa Fluor-568-conjugated goat anti-mouse secondary antibody (Thermo Fisher Scientific Cat#A11031, RRID:AB_144696) at a concentration of 1:200 was applied for 1 h. The cells were stained with DAPI prior to imaging.

High-content imaging was performed on the stained cardiomyocytes using an Operetta CLS High Content Analysis System. These images were processed using CellProfiler [38] using a cellular segmentation algorithm developed previously and validated to within 5% of two independent manual segmentations [39], [40]. Median cell area was used as a representative measure of the cell population in each well, and cells with undetectable cytoplasm were not counted.

### Gene Expression and Bulk RNA Sequencing Analyses

Following the same treatment protocol as described above. RNA extraction and sequencing were performed by Genewiz, Inc. using an Illumina® HiSeq® system. Principal component analysis (PCA) and differential gene expression were conducted using the **DESeq2** package in R [41]. Gene Ontology (GO) enrichment analysis was performed with the **ClusterProfiler** package in R [42]. Transcription factor activity inference was performed using **decoupleR**, with a Univariate Linear Model and CollecTRI regulons, averaging rank across all [43]. To visualize the gene expression of HCM and PI3K-AKT pathways, we used PathView package in R [44]. RNA sequencing data were deposited at GEO (accession # GSE343779).

### Retrospective Clinical Analysis

To examine whether drugs were correlated with a reduction in cardiac adverse events associated with hypertrophy and cardiomyopathy, we analyzed data from the FDA’s Adverse Event Reporting System using the AERSMine tool [45]. We included four different adverse events, including Heart Failure, Ventricular Arrythmias And Cardiac Arrest, Left Ventricular Hypertrophy, and Hypertrophic Cardiomyopathy. The Adverse Event Reporting System is a multicohort database containing 20 million reports of adverse events from healthcare providers and consumers across the United States and across multiple demographics (sex, age, ethnic). The report timeline spans from 2012-2024.

This retrospective study is exempt from informed consent. The data reviewed are a secondary analysis of existing data, do not involve intervention or interaction with human subjects, and are deidentified per the deidentification standard defined in Section §164.514(a) of the HIPAA Privacy Rule. The process by which the data are deidentified is attested to through a formal determination by a qualified expert as defined in Section §164.514(b)(1) of the HIPAA Privacy Rule. This formal determination by a qualified expert was refreshed on December 2020.

Electronic medical records from the University of Virginia Health System were queried to identify patients who were prescribed oral sirolimus, everolimus, or tacrolimus at the time of transthoracic echocardiography performed between 2021 and 2026. Deidentified data were analyzed for left ventricular posterior wall thickness at end-diastole (LVPWd), interventricular septal thickness at end-diastole (IVSd), and combined left ventricular wall thickness, calculated as LVPWd + IVSd. The mTOR inhibitor group included 70 patients (sirolimus, *n* = 45; everolimus, *n* = 25), and the tacrolimus group included 939 patients.

### Statistics

Statistical analyses were performed in Prism 10 (GraphPad) for *in vivo* murine experiments. The presence of statistical outliers was assessed by a ROUT test (2%), and identified outliers were excluded from analysis. Statistical tests were performed with two-tailed analysis. Comparisons between two groups were conducted by Welch’s or Student’s t-test, while comparisons between more than two groups were made by one-way analysis of variance or mixed-effects analysis. Sidak’s multiple comparisons test was performed as post-hoc analysis where appropriate. Mixed effects analysis was used for paired data with more than two groups. A p-value of less than 0.05 was considered statistically significant. Data were represented as mean ± the standard error of the mean. *Ex vivo* image analysis was performed by averaging 3-5 images per mouse across a minimum of three cardiac sections. Data points shown for imaging of cardiac sections are representative of the average across a mouse, with each data point representative of individual mice. Raw data used in the Figures are freely available on figshare at: https://figshare.com/s/eb80b19f903b996993df (Private link for reviewers, will change to public link on acceptance; permanent doi:10.6084/m9.figshare.33249696).

## Results

### MYPBC3^W1082*/W1082*^ mice develop progressive cardiac hypertrophy and fibrosis consistent with human MYBPC3 W1078* truncation

In a patient family at the University of Virginia, we identified a truncating *MYBPC3* mutation due to a premature stop codon at W1078 (human MYBPC3), which was associated with hypertrophic cardiomyopathy (HCM) and sudden cardiac arrest (**Figure 1A,B**). Previous mouse models of *MYBPC3*-associated hypertrophic cardiomyopathy have largely relied on complete genetic ablation of the protein [46], [47], [48]. A premature stop codon at W1078 in the human protein corresponds to W1082 in the murine ortholog (**Figure 1C,E**). To model this mutation *in vivo*, CRISPR/Cas9 genome editing was used to introduce a premature stop codon (TGA) in place of the native tryptophan codon (TGG) at position 1082 in the mouse *Mybpc3* gene (**Figure 1D**). This model is referred to as MYBPC3^W1082*/W1082*^ in the homozygous state.

**Figure 1.**
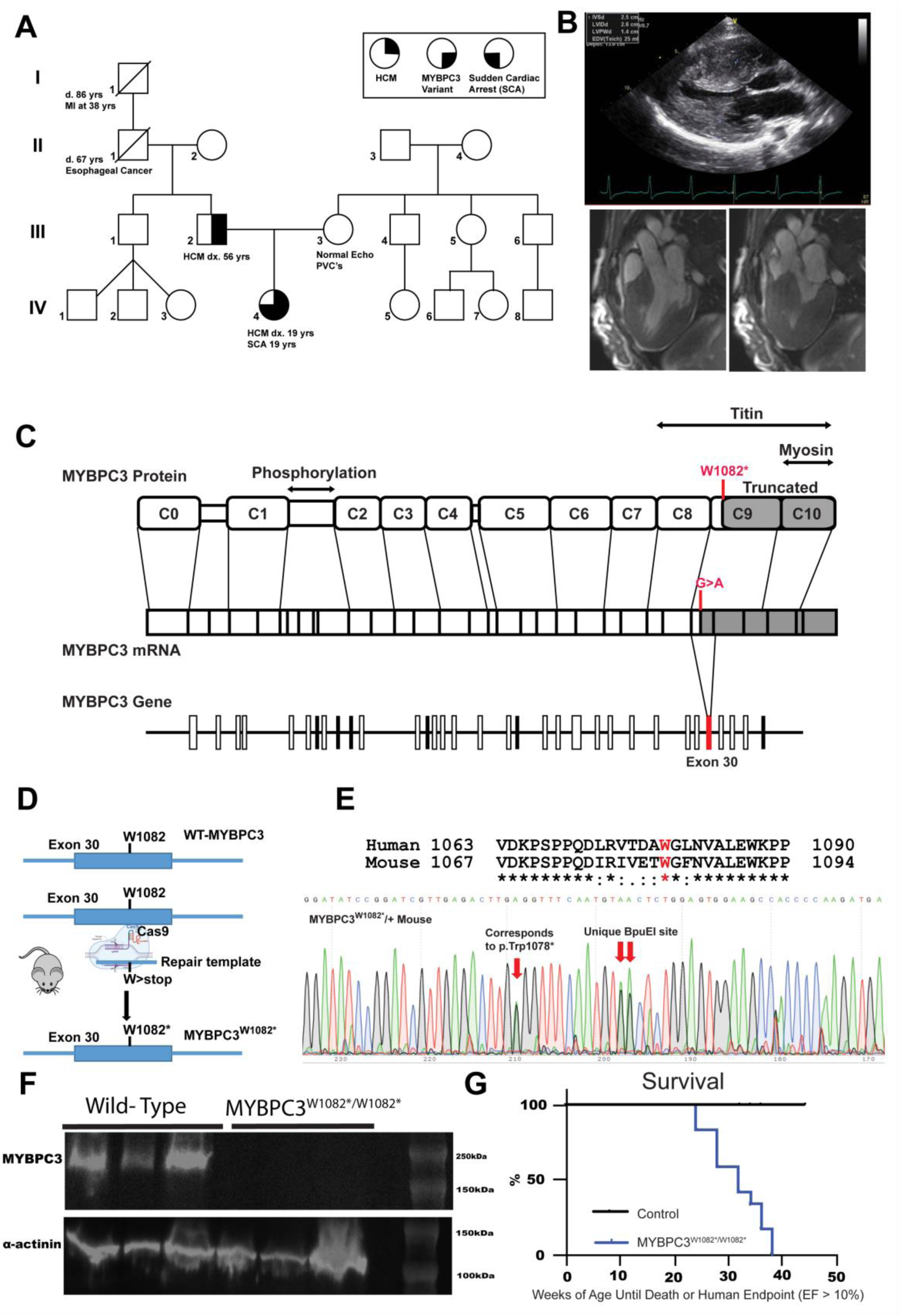
A truncating MYBPC3 variant segregates with early-onset HCM is modeled in vivo. **A)** Pedigree of a multigenerational family with hypertrophic cardiomyopathy (HCM) and sudden cardiac arrest (SCA). Filled symbols denote HCM; half-filled symbols denote variant carriers/SCA. The proband (IV-4, arrow) presented with HCM and SCA at 19 years. Her father (III-2) developed HCM at 56 years. Selected comorbidities are annotated. **B)** Cardiac imaging from the proband demonstrating HCM, with asymmetric septal hypertrophy on echocardiography (top) and concordant structural abnormalities on cardiac magnetic resonance imaging (bottom). **C)** Domain map of MYBPC3 and gene structure highlighting a truncating variant (W1082*) in exon 30. The variant predicts loss of C-terminal domains (C9–C10) required for sarcomeric anchoring to myosin and titin. **D)** CRISPR/Cas9 knock-in strategy introducing the orthologous W1082* (G>A) stop variant into murine Mybpc3. **E)** Allele validation. Top: conservation of the affected residue across species. Bottom: Sanger sequencing confirming the targeted substitution and engineered restriction site. **F)** Western blot for MYBPC3 in whole heart lysate of wild-type and MYBPC3^W1082*/W1082*^ demonstrates no detectable MYBPC3 protein. Loading control of alpha-actinin is shown. **G)** Survival curve of mixed sex wild-type and MYBPC3^W1082*^ mice demonstrate median age of survival of 28 weeks for MYBPC3^W1082*/W1082*^ mice (p=0.006) (WT-N=8, MYBPC3^W1082*/W1082*^-N=12). *P-value determined by Mantel-Cox (G)*.

RNA sequencing of isolated cardiomyocytes demonstrated a significant reduction in *Mybpc3* transcript levels in MYBPC3^W1082*/W1082*^ mice compared with wild-type littermate controls (**Figure 5B**). Consistent with this finding, western blot analysis revealed no detectable MYBPC3 protein in homozygous mutant hearts relative to wild-type littermates (**Figure 1F**).

To evaluate progression of left ventricular dysfunction, MYBPC3^W1082*/W1082*^ mice underwent serial echocardiographic assessment beginning at 8 weeks of age. MYBPC3^W1082*/W1082*^ had significantly reduced ejection fraction as early at 8 weeks of age which declined over time with a median ejection fraction less than 10% by 28 weeks of age. Mutant animals demonstrated significantly reduced survival compared with wild-type controls, with a median survival of approximately 28 weeks (p = 0.0006, Mantel-Cox test; **Figure 1G)**. Survival endpoints included spontaneous death or reaching a humane endpoint defined by severe cardiac dysfunction (ejection fraction <10%), requiring euthanasia.

MYBPC3^W1082*/W1082*^ mice exhibited no developmental abnormalities, as assessed by body weight, kidney, liver, and lung mass, and tibial length in both male and female animals (**Supplementary Figure 1A,B)**. In contrast, MYBPC3^W1082*/W1082*^ mice displayed a marked reduction in ejection fraction (∼49.8% vs. ∼22.8%, p = <0.0001, Student’s t-test; **Figure 2A,B)** accompanied by increased left ventricular wall thickness as measured by echocardiography compared with wild-type littermate controls (septal wall + posterior wall) (∼1.5 mm vs. ∼2.3 mm, p = <0.0001, Student’s t-test; **Figure 2C)**, consistent with cardiac dysfunction associated with HCM. The reduction in ejection fraction was associated with significant increases in both end-systolic and end-diastolic volumes at 12 weeks of age in both male and female mice (**Supplementary Figure 1C,D)**.

**Figure 2:**
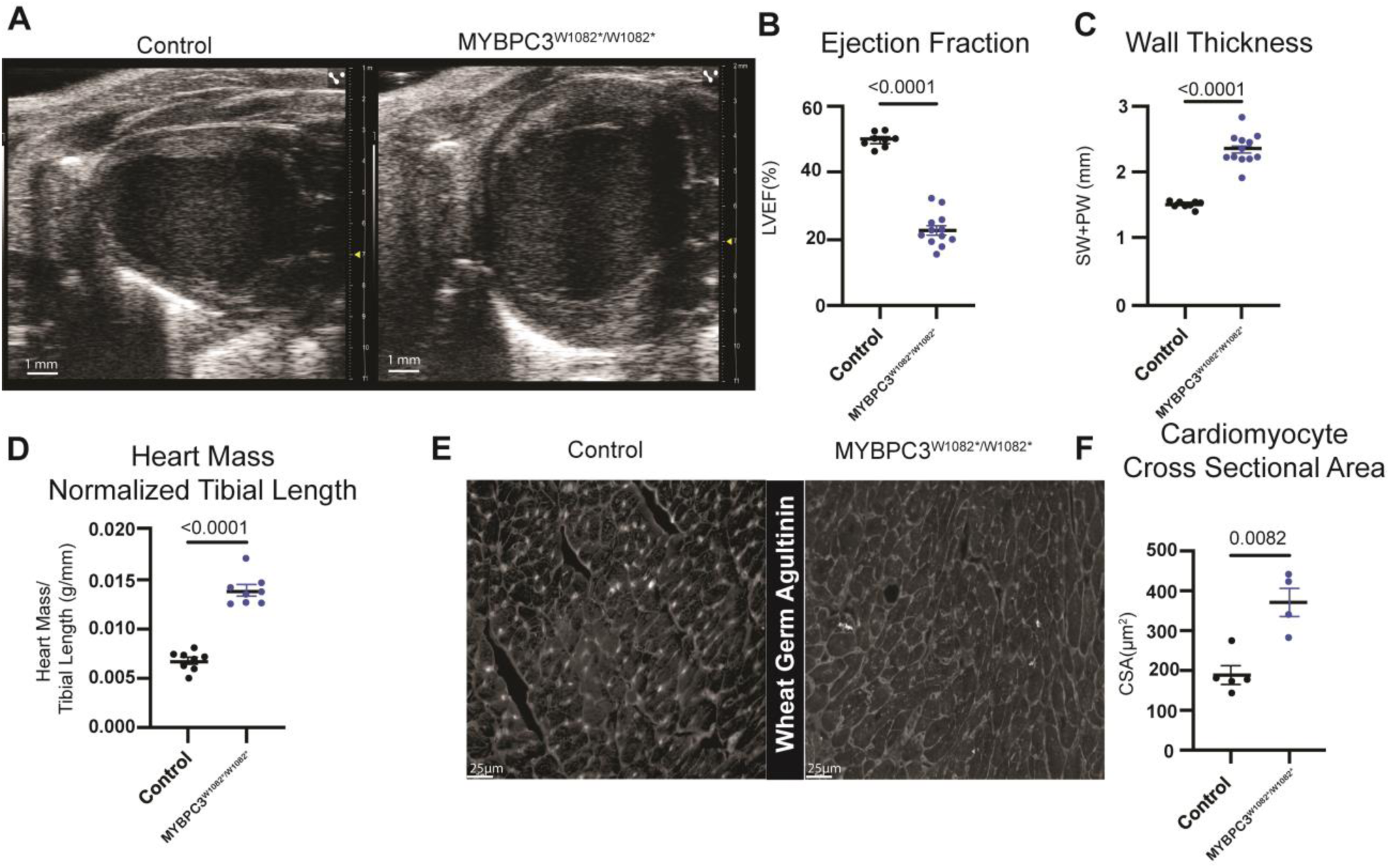
Characterization of a novel murine model of HCM harboring MYBPC3^W1082^*^/W1082^* truncation. **A)** Representative image of B-mode echocardiography of wild-type (left) and MYBPC3^W1082*/W1082*^ (right). **B)** Ejection fraction of wild-type and MYBPC3^W1082*/W1082*^ mice at 8-12 weeks of age is significantly reduced as measured by B-mode echocardiography. **C)** Wall thickness (SW+PW) of wild-type and MYBPC3^W1082*/W1082*^ mice at 8-12 weeks of age is significantly increased as measured by M-mode echocardiography. **D)** Heart mass normalized to tibial length of mixed sex wild-type and MYBPC3^W1082*/W1082*^ mice at 12 weeks of age. Cardiomyocyte cross sectional area **(E)** representative images and **(F)** quantitation as measured by wheat germ agglutinin staining in wild-type and MYBPC3^W1082*/W1082*^ mice at 12 weeks of age. *P-value determined by Student’s t-test (B,C,D), and Welch’s t-test (E). Each individual data point is representative of one animal with data demonstrated as mean* ± *standard error of the mean*.

Histopathological analysis further confirmed an HCM phenotype in MYBPC3^W1082*/W1082*^ mice. MYBPC3^W1082*/W1082*^ animals exhibited a significant increase in heart mass normalized to tibial length (0.007 g/mm vs. ∼0.014 g/mm, p = <0.0001, Student’s t-test; **Figure 2D)**, enlarged cardiomyocyte cross-sectional area measured by wheat germ agglutinin (WGA) staining (∼190 μm^2^ vs. ∼371 μm^2^, p = 0.0082, Welch’s t-test; **Figure 2E,F)**, and sarcomere disorganization evident on hematoxylin and eosin staining (**Supplementary Figure 1E)**. Additionally, MYBPC3^W1082*/W1082*^ hearts displayed significantly increased intramyocardial fibrosis compared with wild-type littermates as assessed by Masson’s trichrome staining (∼1.4% vs. ∼6.5%, p = 0.0489, Welch’s t-test; **Supplementary Figure 1F,G)**.

Collectively, these results establish a novel CRISPR-engineered mouse model that recapitulates a human truncating *MYBPC3* mutation. MYBPC3^W1082*/W1082*^ mice reproduce key features of hypertrophic cardiomyopathy, including ventricular hypertrophy, cardiomyocyte enlargement, sarcomere disorganization, myocardial fibrosis, progressive cardiac dysfunction, and reduced survival. This model therefore provides a clinically relevant platform for investigating the mechanisms and therapeutic targeting of *MYBPC3*-associated HCM.

### A MYBPC3 mutation-specific signaling network model predicts increased cardiomyocyte cell area

To identify the intracellular signaling pathways that drive MYBPC3-associated HCM, we expanded a previously validated logic-based differential equation signaling network model of inherited cardiomyopathies [29]. The base model was curated from experimental literature to encompass 41 nodes representing proteins and mRNAs, connected by 90 regulatory edges. This model exhibited an overall validation accuracy of 83.8% across 160 experimental observations not used to develop the model, spanning HCM, DCM, pressure overload, and volume overload contexts (HCM: 90%; DCM: 75%) [29].

To represent MYBPC3-associated disease, we incorporated a full-length cMyBP-C node, literature-curated regulatory interactions between cMyBP-C and calcium handling pathways, and a MYBPC3^W1082*^ mutation node corresponding to the murine ortholog of the patient-specific MYBPC3^W1078*^ truncation mutation (**Figure 3A**). The model predicts loss of functional cMyBP-C increases myofilament calcium sensitivity while reducing calcium transient amplitude. These alterations propagate through calcium-dependent signaling pathways, resulting in activation of CaMK, NFAT, MEF2, PKG, PKD, and PI3K-AKT signaling. To evaluate the predictive capability of the expanded network, model predictions were compared against independent MYBPC3-specific experimental observations that were not used during model construction. These validation datasets included genetic, pharmacologic, and biomechanical perturbations reported in MYBPC3-associated cardiomyopathy studies. The model correctly reproduced the direction of change for the majority of measured signaling and phenotypic endpoints (**Figure 3B**), supporting its ability to capture key mechanisms downstream of MYBPC3 truncation.

**Figure 3.**
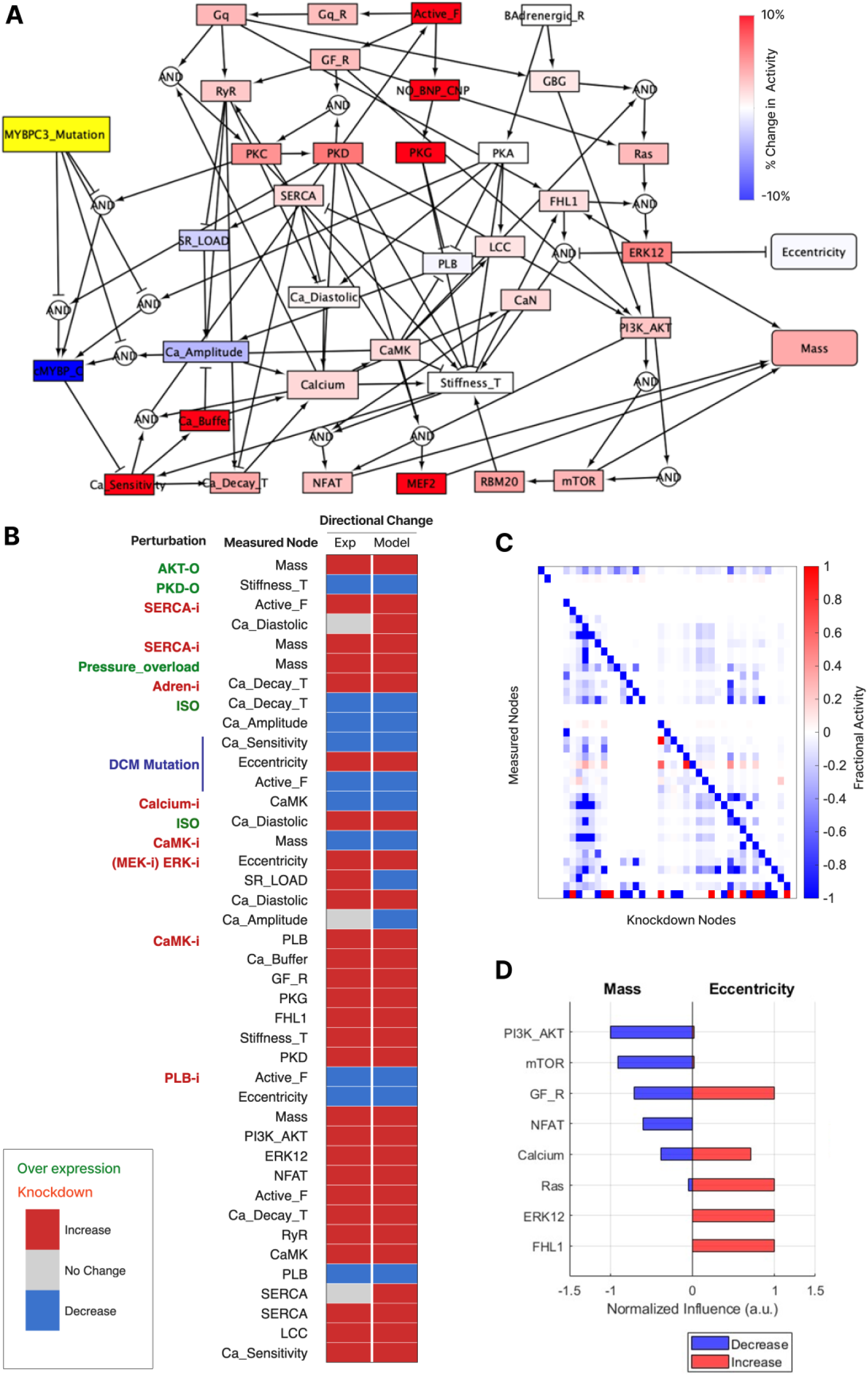
Signaling network model predicts mediators of cardiomyocyte hypertrophy caused by MYBPC3^W1082^* mutation. **(A)** Schematic of the logic-based signaling network model used to simulate the effects of MYBPC3^W1082^* mutation on cardiomyocyte signaling. Nodes represent signaling molecules or cellular processes, and edges represent activating or inhibitory interactions. Node colors indicate the semi-quantitative change in activity in response to MYBPC3 mutation compared to wildtype (red, increased activity; blue, decreased activity). Model outputs representing hypertrophic phenotypes include cardiomyocyte mass and eccentricity. **(B)** Validation of the MYBPC3-expanded signaling network model against independent MYBPC3-specific experimental observations not used for model construction. Experimental observations (Exp) are compared with corresponding model predictions (Model) across literature-derived perturbations and measured signaling or phenotypic endpoints. Red indicates increased activity, blue indicates decreased activity, and gray indicates no significant change. **(C)** Comprehensive virtual knockdown screen evaluating the influence of each network node on the activity of all other nodes. Each column represents a simulated knockdown of the specified node, and each row represents the resulting change in the measured node activity. **(D)** Top five most influential signaling nodes regulating cardiac mass or eccentricity, identified from the virtual knockdown analysis. Bars indicate the normalized influence of each node on mass (blue bars) or eccentricity (red bars).

At steady state, the MYBPC3^W1082*^ mutation was predicted to increase cardiomyocyte mass while reducing eccentricity, consistent with the hypertrophic phenotype observed in the MYBPC3^W1078*/+^ patient and the enlarged cardiomyocytes observed in MYBPC3^W1082*/W1082*^ mice (**Figures 1 and 2E**). Virtual knockdown screen identified PI3K-AKT, mTOR, growth factor receptor (GF_R), NFAT, and calcium signaling as the strongest positive regulators of cardiomyocyte mass, whereas ERK12, FHL1, Ras, and calcium-dependent pathways exerted the greatest influence on eccentricity (**Figure 3D**). These results suggest that activation of PI3K-AKT-mTOR signaling downstream of altered calcium handling is a central mechanism linking MYBPC3 truncation to pathological hypertrophic remodeling.

### Virtual knockdown screen identifies PI3K/AKT-mTOR signaling as the key regulator of MYBPC3-associated cardiomyocyte hypertrophy

To systematically prioritize therapeutic targets, we performed a virtual knockdown screen in which the activity of each node in the network was individually set to zero and the propagating effects on downstream nodes were quantified relative to MYBPC3^W1082^* mutation alone (**Figure 3B**). Network influence was defined as the number of network nodes exhibiting a ≥15% change in activity following knockdown. Highly influential nodes were then used to infer mechanisms underlying mutation-specific disease responses.

This unbiased analysis predicted that PI3K/AKT and mTOR are the most influential pathways controlling cardiomyocyte mass in the context of MYBPC3 mutation context. Virtual knockdown of PI3K, AKT, or mTOR nodes each produced the largest simulated reductions in the cardiac mass output node (**Figure 3C**). In contrast, Ras and ERK1/2 signaling emerged as the dominant regulators of cardiac eccentricity, with the model simulations predicting that their inhibition would promote increased elongation without a corresponding change in cell area (**Figure 3C**). Together, these results suggest that cardiomyocyte size and shape are regulated by partially separable signaling axes downstream of MYBPC3^W1082^* mutation, and that targeting PI3K/AKT and mTOR pathways may selectively attenuate pathological hypertrophy.

### Rapamycin reduces cardiomyocyte hypertrophy in a pharmacologic model of HCM

To translate model predictions into potential candidate therapeutics, we expanded the network model with drug-target interactions for influential protein nodes and then performed a virtual drug screen for drugs that modulate cardiac mass and eccentricity (**Figure 4A**). This virtual screen predicted that in the context of MYBPC3 mutation, the most anti-hypertrophic drugs are inhibitors targeting mTOR (sirolimus) or PI3K (alpelisib, copanlisib, duvelisib, idelalisib).

**Figure 4.**
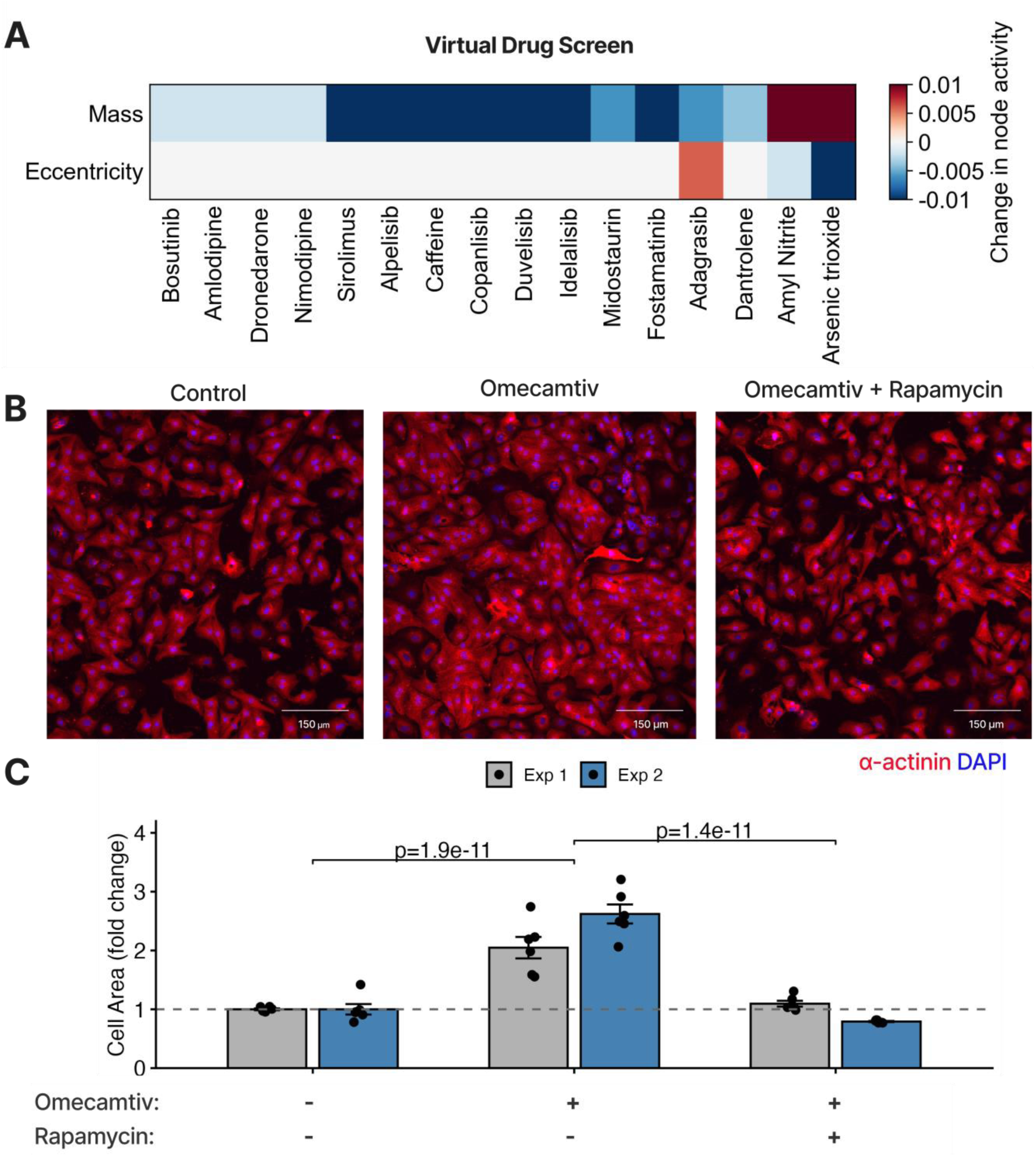
Network-based virtual drug screen in context of MYBPC3 mutation predicts anti-hypertrophic role of sirolimus (rapamycin), validated experimentally in neonatal rat cardiomyocytes. (A) Model-predicted effects of the top 16 candidate drugs on cardiomyocyte phenotypes. Heatmap shows the simulated change in cardiac mass and eccentricity following drug treatment, compared to MYBPC3 mutation alone. (B) Representative immunofluorescence images of NRCMs treated with 10 µM omecamtiv and/or 10 µM rapamycin for 48 h, stained with α-actinin (red) and DAPI (blue). (C) Quantification of NRCM cell area across two independent cell isolations, 6 replicate wells per condition (Exp 1 and Exp 2). Data represent mean ± SEM with individual biological replicates shown. Statistical significance was assessed using a linear mixed-effects model (condition as fixed effect, experiment as random intercept); *p < 0.05, **p < 0.01.

As an in vitro experimental system relevant to hypercontraction-induced hypertrophy characteristic of HCM, we established that the myosin activator omecamtiv mecarbil induces hypertrophy of neonatal rat cardiomyocytes (NRCMs) (p = 0.0003 relative to vehicle treated controls) (**Figure 4B**). Images were acquired by high-content immunofluorescence microscopy of α-actinin and DAPI staining and automated cell segmentation. We then used this system to experimentally validate the model prediction that rapamycin would suppress cardiomyocyte hypertrophy. Treatment with rapamycin significantly attenuated omecamtiv-induced cardiomyocyte hypertrophy (> 2-fold reduction and p < 0.0001) (**Figure 4C**), consistent with model predictions of mTOR pathway selectivity for cardiac mass.

**Figure 5.**
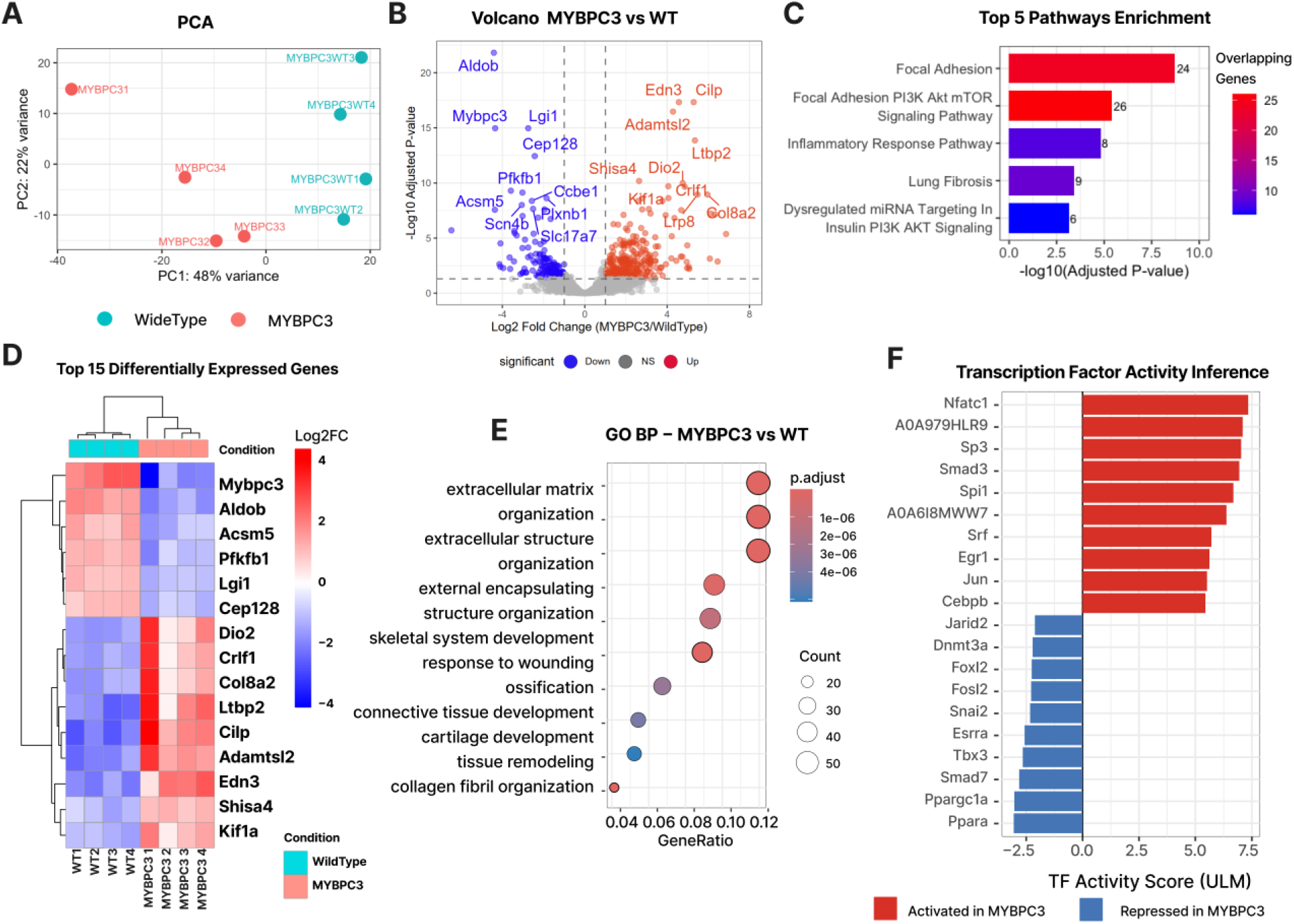
MYBPC3^W1082^* mutant cardiomyocytes exhibit signatures of PI3K/AKT-mTOR signaling and extracellular matrix-related genes. **(A)** Principal component analysis (PCA) of bulk RNA-seq samples showing separation between MYBPC3 mutant and wild-type (WT) cardiomyocytes based on global gene expression profiles. Percent variance explained by each principal component is indicated on the axes. **(B)** Volcano plot of differential gene expression between MYBPC3^W1082*/W2082*^ mutant and WT cardiomyocytes. Significantly upregulated genes are shown in red and downregulated genes in blue, with the top 10 most significant genes labeled. **(C)** Top enriched pathways identified from differentially expressed genes, ranked by – log₁₀(adjusted p-value). Bar color indicates the number of overlapping genes contributing to each pathway. **(D)** Heatmap of the top 15 differentially expressed genes comparing MYBPC3^W1082^* mutant and WT cardiomyocytes, illustrating distinct transcriptional patterns between conditions. **(E)** Gene Ontology (GO) enrichment analysis of biological processes associated with differentially expressed genes. Bubble size indicates gene count and color represents adjusted p-value. **(F)** Transcription factor activity inference identifying the top 10 activated and top 10 repressed transcription factors in MYBPC3^W1082^* mutant cardiomyocytes. Red bars indicate increased predicted increased TF activity, while blue bars indicate repressed TF activity relative to WT.

### Transcriptomic signatures of MYBPC3^W1078*^ cardiomyocytes independently validate PI3K/AKT-mTOR network model predictions

To identify potential mechanisms by which MYBPC3 mutation induces HCM, we performed bulk RNA sequencing of isolated cardiomyocytes from 8–12-week-old male MYBPC3^W1082^*^/W1082*^ mutant mice and wild-type littermate controls. Principal component analysis (PCA) demonstrated clear separation between MYBPC3 mutant and wild-type samples along PC1 (accounting for 48% of total variance), confirming robust condition-specific transcriptional signatures (**Figure 5A**). Differential expression analysis identified total 448 differentially expressed genes in MYBPC3^W1082^* cardiomyocytes (adjusted p < 0.05, |log2FC| >1), with Mybpc3 as the top differentially downregulated gene in MYBPC3^W1082^*^/W1082*^ cardiomyocytes (**Figure 5B and 5D),** which was confirmed by absence of MYBPC3 protein in isolated cardiomyocytes (**Figure 1F**). Gene set enrichment analyses revealed significant overrepresentation of differentially expressed genes associated with PI3K/AKT–mTOR pathways (**Figure 5C**), further validating the network model predictions of a key role for PI3K/mTOR signaling.

To identify upstream transcriptional regulators driving the observed changes in gene expression, we applied decoupleR with 1114 candidate transcription factors (TFs). The top TFs predicted to have increased activity in MYBPC3^W1082*/W1082*^ cardiomyocytes were NFATc1, A0A979HLR9 (Sp1), Sp3, and Smad3, representing all established regulators of hypertrophic gene programs **(Figure 5F)**. TFs with predicted increased activity (**Figure 5G**) regulate highly differentially up-regulated genes, including for NFATc1 (Myh7, Il16), Sp1 (Nppa), and Smad3 (Acta1) [49], [50], [51], [52] (**Supplementary Figure 2**). Conversely, the most repressed TFs included Ppara, Ppargc1a, Smad7, and Tbx3, consistent with metabolic reprogramming and imaged counter-regulatory signaling in pathological hypertrophy [53], [54], [55], [56], [57] (**Figure 5G; Supplementary Figure 2**). Repressed TFs were associated with highly down-regulated genes involved primarily in metabolism rather than cardiac hypertrophy including, Ppara (Slc22a3) and Ppargc1a (Aldob) (**Figure 5G; Supplementary Figure 2**). Together, these computational predictions, cellular validations, and transcriptome profiles converge to identify PI3K/AKT–mTOR signaling as a central driver of pathological cardiomyocyte growth in MYBPC3 mutation-associated HCM and highlight rapamycin as a candidate therapeutic to mitigate disease-associated hypertrophy.

### Rapamycin attenuates cardiac hypertrophy in MYBPC3W1082* mice

To test the therapeutic efficacy of mTOR inhibition *in vivo*, we administered Rapamycin (2 mg/kg/day, i.p.) or vehicle to homozygous MYBPC3^W1082*/W1082^* mutation mice for 28 days, with echocardiographic assessment at baseline, 2 weeks, and 4 weeks, and organ harvest at study endpoint (**Figure 6A**). Rapamycin treatment did not affect body mass in mice 6-10 weeks of age (p=0.3295, **Supplement Figure 3).**

**Figure 6:**
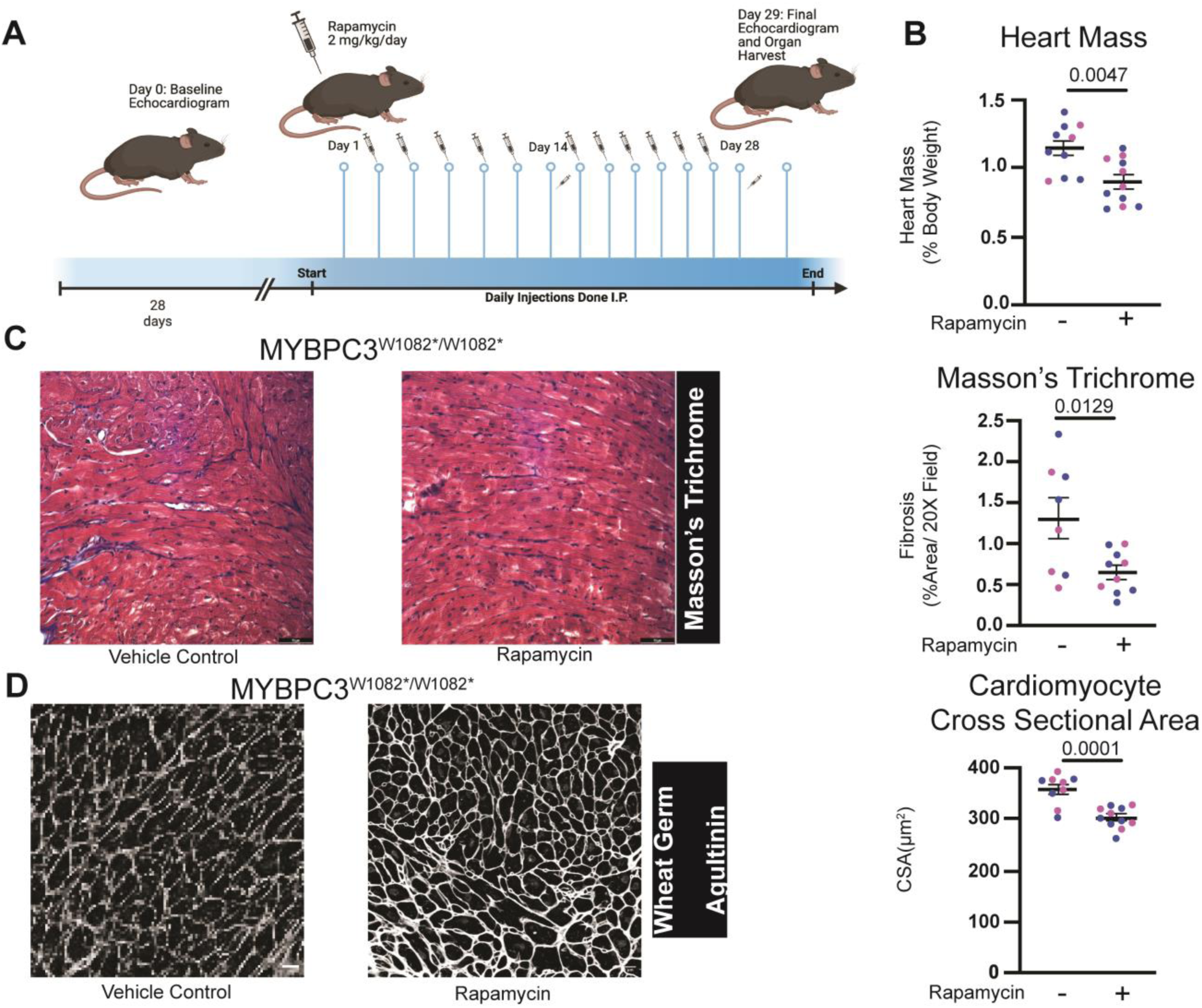
Rapamycin attenuates cardiac hypertrophy and fibrosis in MYBPC3^W1082*/W1082*^ mice. **(A)** Schematic describing rapamycin (2mg/kg/day) administration over 28 days in MYBPC3^W1082*/W1082*^ mice. **(B)** Heart mass normalized to body weight of vehicle control or rapamycin administered MYBPC3^W1082*/W1082*^ mice in a mixed sex cohort (male = blue circle, female = pink circle). **(C)** Interstitial fibrosis as measured by Masson’s trichrome (representative images left, quantification right) in vehicle control or rapamycin administered MYBPC3^W1082*/W1082*^ mice. **(D)** Cardiomyocyte cross sectional area representative images (left) and quantitation (right) as measured by wheat germ agglutinin staining in vehicle control or rapamycin administered MYBPC3^W1082*/W1082*^ mice. P-value determined by Student’s t-test. *Each individual data point is representative of one animal with data demonstrated as mean* ± *standard error of the mean*.

Rapamycin treatment produced a significant reduction in cardiac mass in MYBPC3^W1082*/W1082^* mutation mice. Heart mass normalized to body weight was significantly lower in rapamycin-treated mutant mice compared to vehicle-treated controls (∼0.90% vs. ∼1.15%, p = 0.0047, Student’s t-test; **Figure 6B).** Heart mass normalized to tibial length was similarly reduced (∼0.012 g/mm vs. ∼0.016 g/mm, p = 0.0084; **Figure 6B**). Wall thickness (SW+PW) (p = 0.7764; **Supplementary Figure 3),** and ejection fraction were not significantly altered by Rapamycin treatment (p = 0.7313; **Supplementary Figure 3**).

Histological analyses confirmed beneficial tissue-level remodeling with Rapamycin treatment. Masson’s trichrome staining revealed a significant reduction in myocardial fibrosis in Rapamycin-treated MYBPC3^W1082*/W1082^* hearts compared to vehicle-treated mutants (∼0.64% vs. ∼1.29% positive area, p = 0.0129, Student’s t-test; **Figure 6C).** Cardiomyocyte cross-sectional area was also significantly reduced following Rapamycin treatment (∼303 µm² vs. ∼358 µm², p = 0.0001; Student’s t-test; **Figure 6D),** demonstrating that mTOR inhibition attenuates pathological cardiomyocyte hypertrophy at the cellular level. Together, these *in vivo* data demonstrate that pharmacological mTOR inhibition with Rapamycin significantly reduces cardiac mass, fibrosis, and cardiomyocyte hypertrophy in MYBPC3^W1082*/W1082*^ mice though it does not acute improve cardiac function.

### Sirolimus (rapamycin) use in patients is associated with reduced incidence of adverse hypertrophic cardiac events

To assess the translational potential of mTOR inhibition for reducing cardiac hypertrophy in humans, we performed a retrospective analysis of the FDA Adverse Event Reporting System (FAERS) using the AERSMine tool [45], examining reports of hypertrophy-related cardiac events from 2012–2024. We compared event frequencies among patients prescribed sirolimus (rapamycin) or everolimus (both mTOR inhibitors) versus tacrolimus, a calcineurin inhibitor commonly used for the same indication of solid organ transplant immunosuppression but with a mechanistically distinct mode of action. The analysis included 263,122 sirolimus-prescribed patients, 68,301 everolimus-prescribed patients, and 173,284 tacrolimus-prescribed patients serving as the reference group.

Tacrolimus-treated patients had a left ventricular hypertrophy reporting rate of 12.9 per 10,000 patients. Both mTOR inhibitors were associated with significantly lower left ventricular hypertrophy reporting rates relative to tacrolimus (**Figure 7**). Sirolimus-treated patients had a reporting rate of 10.5 per 10,000 (ROR 0.81, 95% CI 0.68–0.97, p = 0.022), and everolimus-treated patients had a rate of 6.2 per 10,000 (ROR 0.48, 95% CI 0.34–0.66, p < 0.0001), representing 19% and 52% lower reporting odds than tacrolimus treatment, respectively. We further examined incidence of reported hypertrophic cardiomyopathy, a related phenotype. The hypertrophic cardiomyopathy reporting rate for tacrolimus was 9.8 per 10,000. Sirolimus-treated patients had a significantly lower hypertrophic cardiomyopathy reporting rate of 7.2 per 10,000 (ROR 0.74, 95% CI 0.60–0.91, p = 0.0037), and everolimus-treated patients had the lowest rate at 2.6 per 10,000 (ROR 0.27, 95% CI 0.17–0.44, p < 0.0001) (**Figure 7**). These correspond to a 26% and 73% reduction in reporting odds relative to tacrolimus, respectively. Compared to patients on tacrolimus, patients on everolimus or sirolimus also exhibited a lower incidence of ventricular arrhythmias, cardiac arrest, or ventricular fibrillation (**Supplementary Figure 4**).

**Figure 7.**
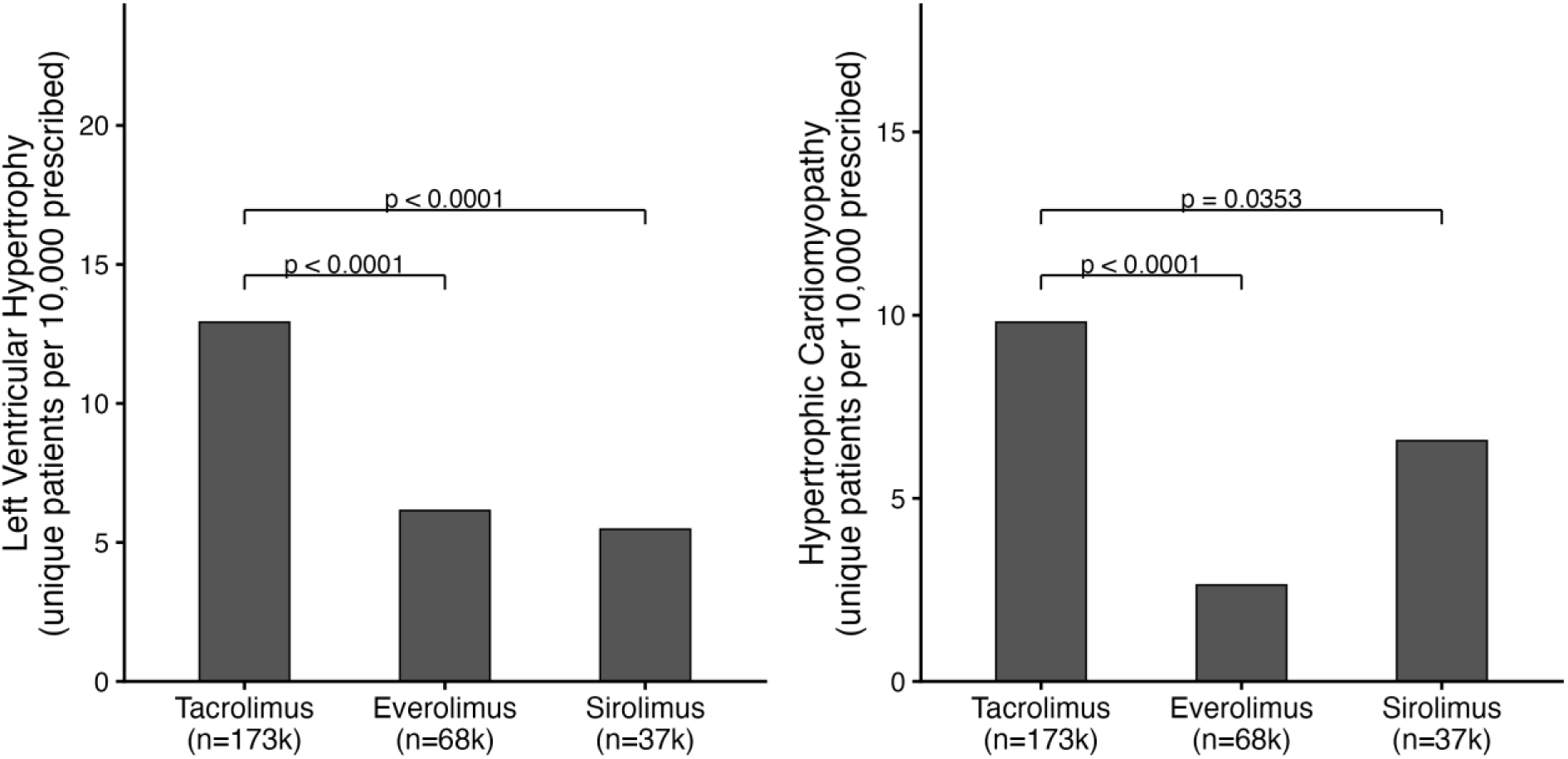
Cardiac hypertrophy adverse event reporting rates in the FDA Adverse Event Reporting System (FAERS) for mTOR inhibitors versus tacrolimus. Unique patients reporting each adverse event per 10,000 patients prescribed, extracted from FAERS via AERSmine. (A) Left ventricular hypertrophy (LVH) and (B) Hypertrophic cardiomyopathy (HCM) reporting rates for tacrolimus (n = 173,284), everolimus (n = 68,301), and sirolimus (n = 263,122). Bars represent the reporting rate per 10,000 unique patients prescribed each drug. Comparison brackets indicate Reporting Odds Ratio (ROR) analysis with Woolf 95% confidence intervals relative to tacrolimus as the reference. P-values are derived from the normal approximation of the log-ROR.

To explore whether chronic mTOR inhibition is associated with differences in cardiac wall thickness in an independent patient population, we queried the University of Virginia electronic medical records system to identify patients prescribed oral sirolimus, everolimus, or tacrolimus at the time of transthoracic echocardiography performed over the preceding five years (2021-2026). Because sirolimus and everolimus share a common mechanism as mTOR inhibitors, these cohorts were combined and compared with tacrolimus-treated patients. The final analysis included 70 patients receiving mTOR inhibitors (45 sirolimus and 25 everolimus) and 939 patients receiving tacrolimus.

Patients treated with sirolimus or everolimus exhibited significantly lower left ventricular posterior wall thickness in diastole (LVPWD) compared with tacrolimus-treated patients (1.01 ± 0.18 cm versus 1.08 ± 0.22 cm, *P* = 0.0042; **Figure 8A, D-G**). Similarly, interventricular septal thickness in diastole (IVSD) was reduced in the mTOR inhibitor group relative to the tacrolimus group (1.04 ± 0.19 cm versus 1.10 ± 0.23 cm, *P* = 0.0104; **Figure 8B, D-G**). To assess overall ventricular wall thickness, LVPWD and IVSD measurements were summed for each patient. The combined wall thickness remained significantly lower in patients receiving sirolimus or everolimus compared with tacrolimus-treated patients (2.05 ± 0.34 cm versus 2.18 ± 0.42 cm, *P* = 0.0034; **Figure 8C**).

**Figure 8:**
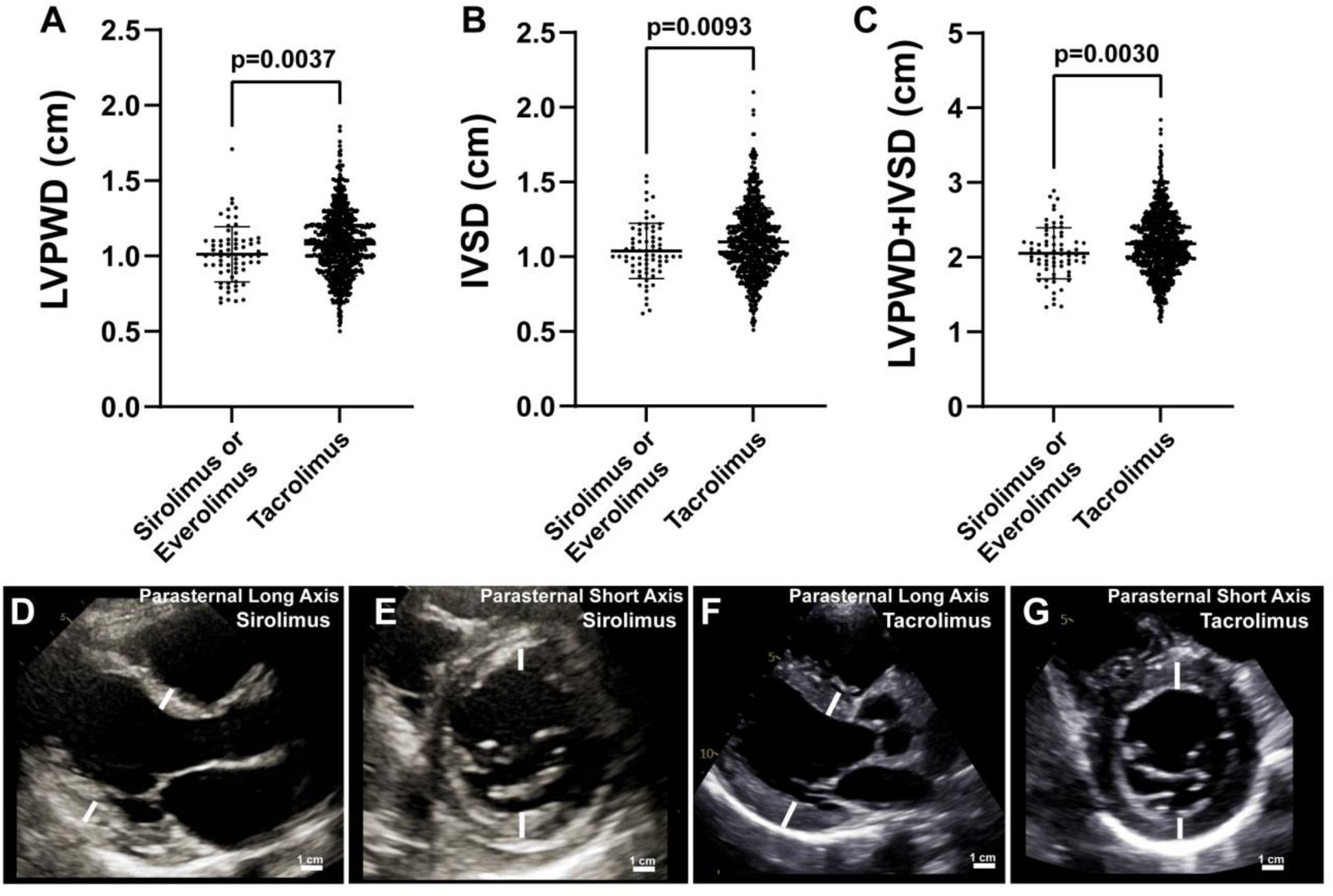
Reduced left ventricular wall thickness in patients receiving mTOR inhibitors compared with tacrolimus. Electronic medical records from the University of Virginia Health System were queried to identify patients prescribed oral sirolimus, everolimus, or tacrolimus at the time of transthoracic echocardiography performed during the preceding five years. Because sirolimus and everolimus share a common mechanism of action as mTOR inhibitors, these cohorts were combined for analysis and compared with tacrolimus-treated patients. Scatter plots depict individual patient measurements with horizontal bars representing mean ± SD. **(A)** Left ventricular posterior wall thickness in diastole (LVPWD). **(B)** Interventricular septal thickness in diastole (IVSD). **(C)** Total left ventricular wall thickness calculated as LVPWD + IVSD. The mTOR inhibitor group (sirolimus, *n* = 45; everolimus, *n* = 25; combined *n* = 70) exhibited significantly lower LVPWD (1.01 ± 0.18 cm vs. 1.08 ± 0.22 cm, *P* = 0.0042), IVSD (1.04 ± 0.19 cm vs. 1.10 ± 0.23 cm, *P* = 0.0104), and combined wall thickness (2.05 ± 0.34 cm vs. 2.18 ± 0.42 cm, *P* = 0.0034) compared with tacrolimus-treated patients (*n* = 939). Statistical significance was determined using unpaired two-tailed *t*-tests. Representative echocardiographic images are shown from patients receiving sirolimus (**D, E**) or tacrolimus (**F, G**). **(D, F)** Parasternal long-axis views and **(E, G)** parasternal short-axis views obtained at the level of the mitral valve. White bars indicate the left ventricular posterior wall and interventricular septum used for thickness measurements. Representative images illustrate the comparatively reduced ventricular wall thickness observed in patients receiving sirolimus relative to tacrolimus.

Although the absolute differences were modest, the findings were consistent across all three echocardiographic measures and were observed despite the substantially larger tacrolimus cohort. The distributions of measurements demonstrated considerable overlap between groups, indicating that the effect size is relatively small at the individual patient level. Nevertheless, the concordant reduction in posterior wall thickness, septal thickness, and total ventricular wall thickness among patients receiving mTOR inhibitors suggests a potential association between mTOR pathway inhibition and reduced myocardial hypertrophic remodeling in humans. These observations are consistent with extensive experimental literature implicating mTOR signaling as a central regulator of cardiomyocyte growth and pathological cardiac hypertrophy and provide clinical evidence that pharmacologic mTOR inhibition may influence ventricular wall architecture in patients receiving long-term immunosuppressive therapy.

These observational data in a real-world patient population provide complementary clinical evidence that mTOR pathway inhibition may confer cardiac protection against pathological hypertrophic remodeling, consistent with our experimental findings across signaling network, cellular, and in vivo models.

## Discussion

In this study, we initially identified a patient with a truncating mutation in MYBPC3 caused by W1078* mutation. We combined a patient-specific mouse model (MYBPC3^W1082*/W1082*^), a mutation-specific signaling network model, transcriptomic profiling, and pharmacological intervention to identify PI3K/AKT–mTOR signaling as a central driver of pathological hypertrophy in MYBPC3-associated HCM. Furthermore, we demonstrate that inhibition of mTOR with rapamycin in this model attenuates cardiac remodeling *in vivo*. These findings advance our understanding of mutation-specific disease mechanisms and mTOR inhibition as a candidate disease-modifying strategy deserving further translational investigation.

A key strength of this work is the use of a precision CRISPR knock-in mouse carrying the exact truncating variant (p.W1082*/W1082*) orthologous to a human HCM-causing MYBPC3^W1078*/+^ mutation identified in a patient family from the University of Virginia. Unlike conventional transgenic overexpression models or complete knockouts, which introduce non-physiological gene dosage and phenotypes that diverge substantially from human disease, ^[26–29]^ our model recapitulates the molecular lesion present in patients. Homozygous (MYBPC3^W1082*/W1082*^) mice developed progressive LV hypertrophy, sarcomeric disarray, interstitial fibrosis, and reduced ejection fraction, encompassing a phenotypic profile consistent with human HCM. This novel murine model provides a well-characterized platform for mechanistic and therapeutic investigation.

The MYBPC3 signaling network yielded a mechanistically interpretable and experimentally validated map of hypertrophic signaling. A virtual knockdown screen with this network model identified PI3K/AKT–mTOR as the dominant regulator of cardiomyocyte mass and Ras/ERK as the dominant regulator of elongation, which suggests MYBPC3 mutation induces partially separable signaling axes for cardiomyocyte size and elongation. This dissociation has potential clinical implications: therapeutic strategies targeting mass without altering cell geometry may be achievable, and could avoid the adverse remodeling (e.g., chamber dilation) associated with broad suppression of hypertrophic signaling. ^[32,33]^ The orthogonal validation of these predictions by RNA-seq in MYBPC3^W1082*/W1082*^ cardiomyocytes further validated the activation of PI3K/mTOR-dependent gene expression, including activation of NFATc1 and Sp1 transcription factors consistent with known hypertrophic programs. ^[17,18]^

The *in vivo* efficacy of Rapamycin in reducing cardiac mass, cardiomyocyte hypertrophy, and fibrosis in MYBPC3^W1082*/W1082*^ mice is an encouraging finding with direct translational relevance. mTOR inhibition has previously been shown to attenuate pressure overload–induced hypertrophy and improve survival in both rodent and feline models of heart failure and hypertrophic cardiomyopathy, and rapamycin analogs are already in widespread clinical use as immunosuppressants in solid organ transplantation, providing an established safety and pharmacokinetic profile [58], [59], [60]. mTOR inhibition has also been shown to reduce autophagy in a distinct Mybpc3-targeted c.772G>A knock-in mouse [61]. Moreover, we here demonstrate that the beneficial effects of Rapamycin treatment in HCM can be linked with both cardiomyocyte hypertrophy and a reduction in interstitial fibrosis, posing a multifactorial role for mTOR in the HCM myocardium. The concordance between our network predictions, *in vitro* NRCM experiments, *in vivo* Rapamycin treatment, and the FAERS population-level data, where sirolimus use was associated with reduced hypertrophy-related events and wall thickness relative to tacrolimus, creates a multi-scale, convergent evidence base that is unusual in preclinical HCM research. Notably, the preservation of EF under Rapamycin treatment compared with baseline treatment contrasts with the contractility risks associated with mavacamten, suggesting that mTOR inhibition may offer a mechanistically complementary therapeutic approach, particularly for non-obstructive HCM or patients with borderline systolic function[62].

Several limitations warrant consideration. First, the 28-day treatment window does not address whether Rapamycin can sustain regression of hypertrophy or prevent disease onset with earlier intervention. This is of note as while MYBPC3^W1082*/W1082*^ have an overt decline in cardiac function as early as 6-weeks of age, in the clinical setting ejection fraction is largely normal in patients. Additionally, the FAERS analysis is observational and subject to confounding by indication, as sirolimus and tacrolimus are prescribed in overlapping but not identical patient populations; these data should be interpreted as hypothesis-generating rather than confirmatory. In support of this data though a real-world clinical observation demonstrated that patients receiving chronic mTOR inhibitor therapy exhibit modest but consistent reductions in ventricular wall thickness compared with patients receiving tacrolimus. Although retrospective and subject to confounding, these findings provide translational support for experimental studies identifying mTOR signaling as a central regulator of cardiomyocyte growth. Finally, while the network model achieved strong overall prediction accuracy, it does not yet incorporate cell-type heterogeneity (e.g., fibroblast–cardiomyocyte crosstalk) or spatial signaling gradients that may influence therapeutic response *in vivo*.

In summary, this work demonstrates the power of a tightly integrated computational-experimental strategy to uncover mutation-specific therapeutic targets in HCM. By anchoring discovery in a clinically derived variant, validating predictions across network modeling, transcriptomics, cell culture, and a relevant mouse model, and corroborating findings with population-level adverse event data, we provide a multi-scale rationale for mTOR inhibition as a disease-modifying strategy in MYBPC3-associated HCM. Future studies should evaluate the efficacy of rapamycin or other mTOR inhibitors in heterozygous (MYBPC3^W1082*/+^) mice, determine optimal treatment timing and duration, and explore whether combination with sarcomere-directed therapies offers additive or synergistic benefit. Ultimately, the integrated framework developed here is broadly applicable to other sarcomeric mutations and may accelerate mutation-tailored drug discovery across the genetic spectrum of HCM.

## Supplementary Figures

**Supplementary Figure 1:**
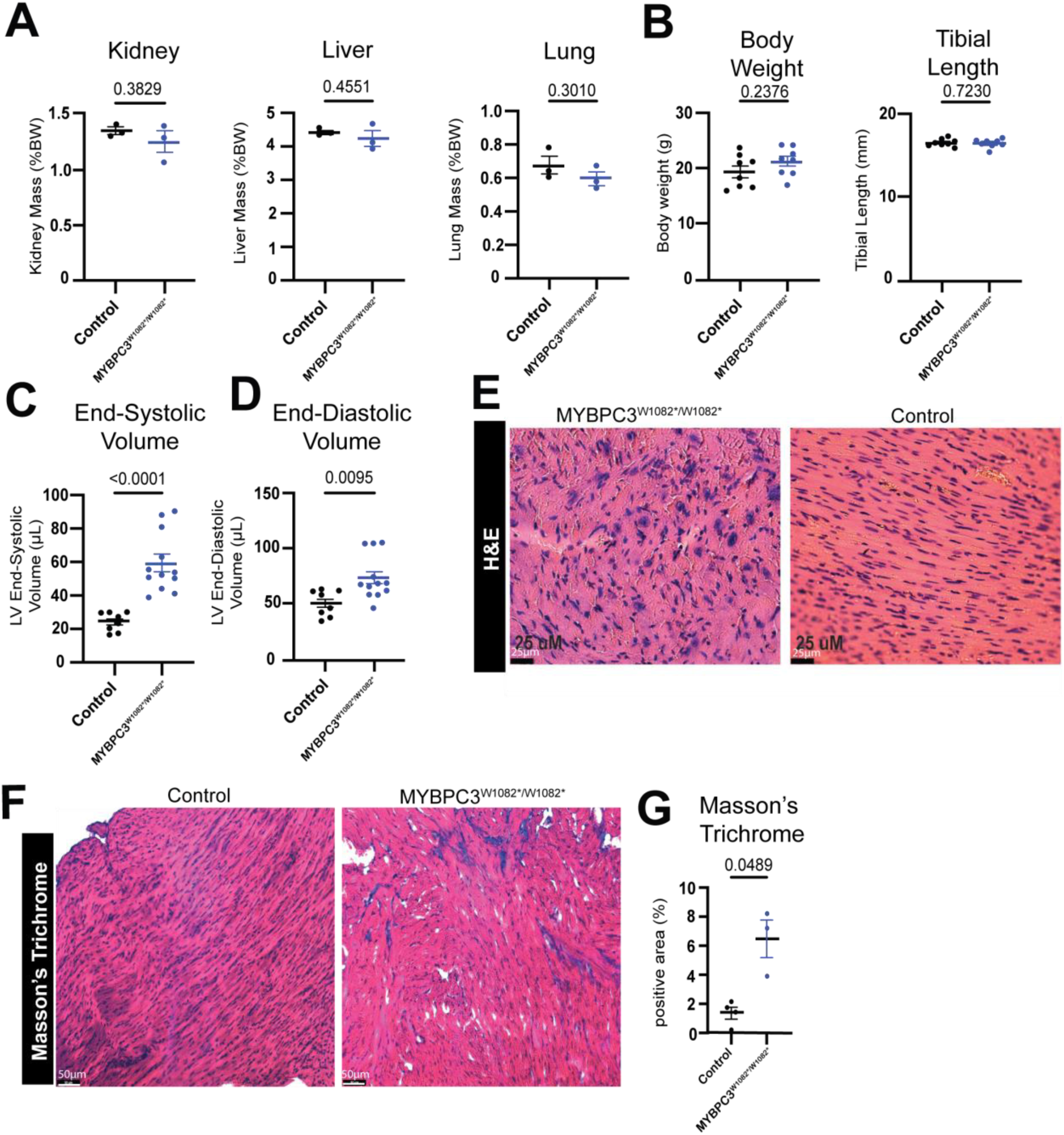
A) Kidney, liver, and lung mass relative to body weight were measured in a mixed sex 12-week old wild-type littermate controls or MYBPC3^W1082*/W1082*^ mice and showed no differences. **B)** Body weight and tibial length as markers of development were measured in 12 week old mixed sex cohort of wild-type littermate controls or MYBPC3^W1082*/W1082*^ mice and showed no significant differences. B-mode echocardiography was used to measure left ventricular **C)** end-systolic and **D)** end-diastolic volume in a mixed sex cohort at 12 weeks of age of wild-type littermate controls or MYBPC3^W1082*/W1082*^ mice demonstrating significant increases in both metrics. **E)** Representative images of H&E in wild-type littermate controls or MYBPC3^W1082*/W1082*^ mice demonstrate gross sarcomere disorganization. Masson’s Trichrome was measured in mixed sex 8-week old control or MYBPC3^W1082*/W1082*^ mice and showed (representative images **F)** a significant increase in fibrosis measured by the **G)** positive trichrome area. *P-value determined by Student’s t-test*.

**Supplementary Figure 2:**
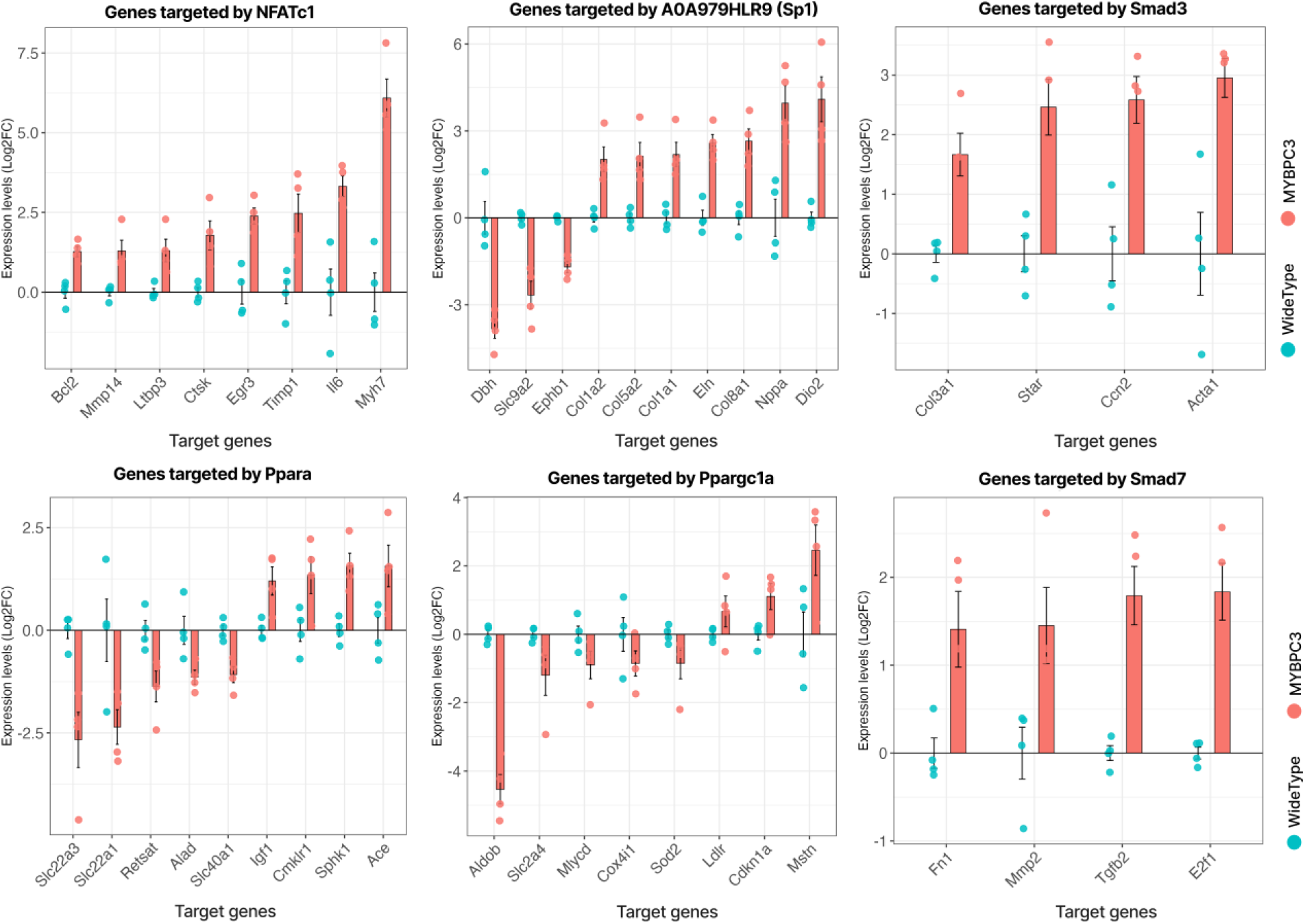
Change in expression of representative target genes regulated by the inferred transcription factors for WT and MYBPC3^W1082^* adult cardiomyocytes. Bars represent mean shown as log₂ fold change expression, with individual data points overlaid for WT and MYBPC3^W1082^* samples.

**Supplementary Figure 3:**
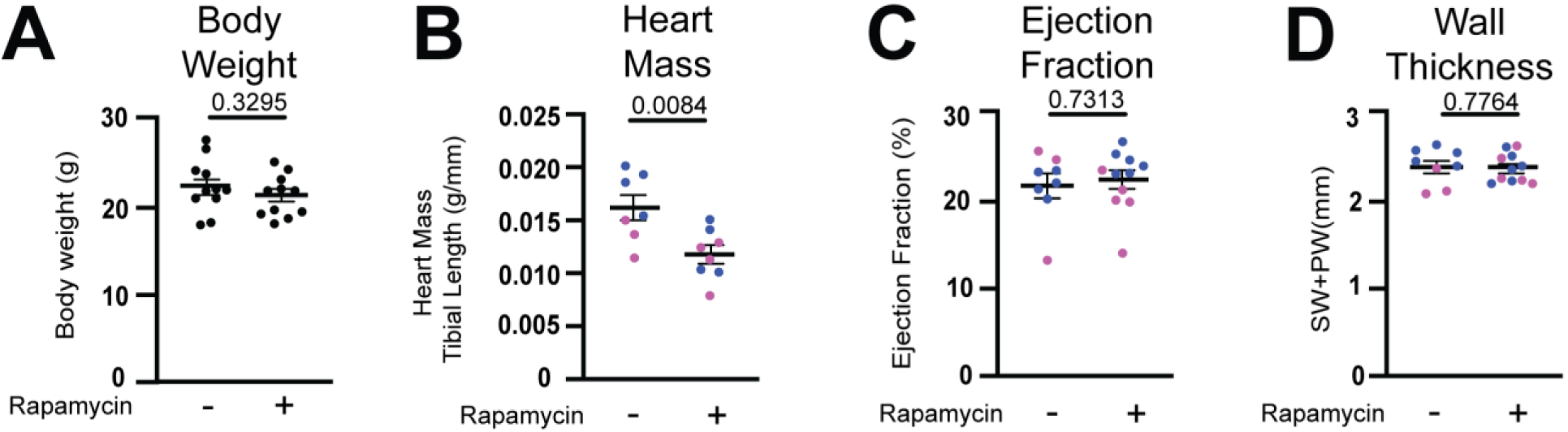
Effects of rapamycin on additional cardiac phenotypes in MYBPC3^W1082*/W1082*^ mice. **A)** Body weight of MYBPC3^W1082*/W1082*^ mice is not altered by daily Rapamycin injection compared with vehicle control. **B)** Heart mass normalized to tibial length was significantly reduced in Rapamycin treated MYBPC3^W1082*/W1082*^ mice compared with vehicle control treatment. B-mode echocardiography was used to measure left ventricular **C)** ejection fraction and showed no significant difference between Rapamycin treated and vehicle control treatment after 4 weeks of injections in MYBPC3^W1082*/W1082*^ mice. M-mode echocardiography was used to measure left ventricular **D)** wall thickness (septal wall + posterior wall) and showed no significant difference between Rapamycin treated and vehicle control treatment after 4 weeks of injections in MYBPC3^W1082*/W1082*^ mice. *P-value determined by Student’s t-test*.

**Supplementary Figure 4.**
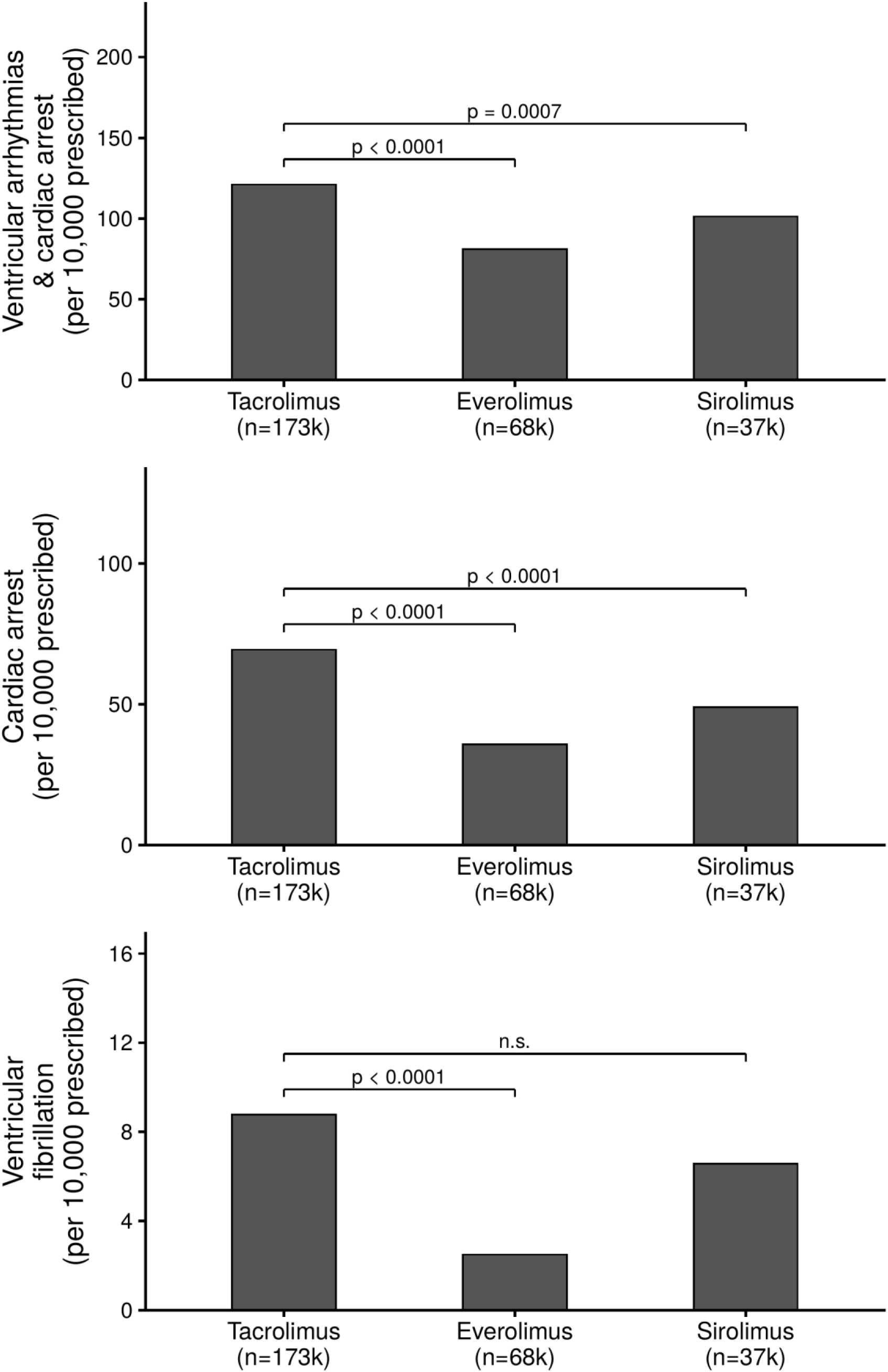
Patients prescribed mTOR inhibitors (sirolimus and everolimus) exhibited lower reporting rates of ventricular arrhythmia/cardiac arrest, cardiac arrest, and ventricular fibrillation in FAERS, compared to calcineurin inhibitor, tacrolimus. Bars show the number of unique patients with a report of each adverse event per 10,000 patients prescribed tacrolimus (n = 173,284), everolimus (n = 68,301), or sirolimus (n = 36,529), based on FAERS data queried via AERSMine. For each adverse event, everolimus and sirolimus were compared to tacrolimus (reference) using a one-sided Fisher’s exact test for a lower reporting rate (protective direction); bracketed values indicate the resulting p-value (n.s., not significant, p ≥ 0.10).

